# Habitat diversification promotes environmental selection in planktonic prokaryotes and ecological drift in microbial eukaryotes

**DOI:** 10.1101/161091

**Authors:** Ramiro Logares, Sylvie V. M. Tesson, Björn Canbäck, Mikael Pontarp, Katarina Hedlund, Karin Rengefors

## Abstract

Whether or not communities of microbial eukaryotes are structured in the same way as prokaryotes is a basic and poorly explored question in ecology. Here we investigated this question in a set of planktonic lake microbiotas in Eastern Antarctica that represent a natural community ecology experiment. Most of the analysed lakes emerged from the sea during the last 6,000 years, giving rise to waterbodies that originally contained marine microbiotas and that subsequently evolved into habitats ranging from freshwater to hypersaline. We show that habitat diversification has promoted environmental selection driven by a salinity gradient in prokaryotes and ecological drift in microeukaryotes. Nevertheless, we detected also a number of microeukaryotes with specific responses to salinity, indicating that albeit minor, environmental selection has had a role in the assembly of their communities. Altogether, we conclude that habitat diversification can promote contrasting responses in planktonic microeukaryotes and prokaryotes belonging to the same communities.

## INTRODUCTION

A fundamental question in ecology is what determines how species are distributed across time and space. According to the framework proposed by Vellend (2010), patterns in the composition of communities can be understood in terms of four main processes: selection, (ecological) drift, dispersal and speciation. Selection represents a deterministic fitness difference between individuals of different species. Ecological drift is associated to stochastic changes in the relative abundance of taxa. Dispersal is the movement of species across space and speciation generates new species. The species composition of a given community is expected to be the product of the latter processes acting locally (e.g. selection or drift) or regionally [dispersal] (Ricklefs 1987). The role of dispersal has been emphasized within the metacommunity concept, which concerns local communities that are linked to each other by the dispersal of multiple and potentially interacting species (Leibold *et al*. 2004). The previous frameworks derive mostly from the study of animals and plants, but lately they have been also applied to microbial communities (Martiny *et al*. 2006; Hanson *et al*. 2012; Lindström & Langenheder 2012;Stegen *et al*. 2013).

Environmental selection (a.k.a species sorting), one of the paradigms within the metacommunity theory, is expected when most species in a metacommunity respond to moderate-high selection under intermediate levels of dispersal (Lindström & Langenheder 2012). This will result in a covariation (or coupling) between community and habitat dissimilarities, and habitats exerting similar environmental selection are expected to contain similar microbial assemblages. Thus, environmental selection is a deterministic force that contrasts with ecological drift, a stochastic process of community assembly that could also include historical effects (Chase 2003). In free-living prokaryotes, environmental selection has been shown to be the prevalent factor structuring communities (Lindström & Langenheder 2012, Pontarp *et al*. 2012) although other studies support neutral or stochastic factors (drift) as having a role in their community assembly (Ofiteru *et al*. 2010). Together with prokaryotes, microbial eukaryotes constitute most microbial communities. Even though microeukaryotes are key components of communities, their study lags behind the study of prokaryotes (Keeling & Del Campo 2017). Consequently, we know very little on whether or not their communities are structured by the same processes as in prokaryotes. Microbial eukaryotes, which have cells that are structurally more complex than prokaryotes, including organelles for different functions [e.g. the vacuole contractile for osmoregulation, peduncles for phagotrophic feeding, chloroplasts for photosynthesis] (Massana & Logares 2013), could present a wide array of responses to environmental heterogeneity, which may differ from prokaryotes.

Here, we explore the question on whether or not prokaryotes and microbial eukaryotes from the same communities are structured by the same processes. We investigate this question in planktonic microbial communities populating lakes of an Antarctic region that can be considered a natural experiment in community ecology. This region (Vestfold Hills) has about 411 km^^2^^ and contains ca. 300 lakes (Laybourn-Parry & Pearce 2007; Cavicchioli 2015; Fig. S1). Due to post-glacial rebound, coastal landmasses emerged from the sea in the Vestfold Hills, leaving behind isolated pockets of water with their corresponding landlocked marine microbiotas (Zwartz *et al*. 1998; Gibson *et al*. 2009). During the last 6,000 years, several lakes gradually evolved from ancestral marine conditions into more saline or less saline environments (Zwartz *et al*. 1998; Gibson *et al*. 2009). Today, the Vestfold Hills contains a mosaic of lakes ranging from freshwater (salinity 0) to hypersaline (salinity 250) with a separation between them which does not exceed 20 km (Fig. S1). The same range of distances separate the lakes and the sea, a potential source of immigrants. Overall, habitat diversification exposed progressively ancestral marine microbiotas to new regimes of environmental selection and opened new niches for immigrant taxa.

A previous study has shown that environmental selection driven by a salinity gradient has been the main process shaping the composition of prokaryotic communities in the region (Logares *et al*. 2013). Salinity is a major factor determining the spatial distribution of prokaryotes (Lozupone & Knight 2007) and salinity differences are also a major barrier for environmental transitions in both prokaryotes and microeukaryotes (Logares *et al*. 2007; Logares *et al*. 2009). Here we explore whether microeukaryotes populating the same lakes in this region are structured, as prokaryotes, by salinity-driven environmental selection or other processes.

In order to compare microeukaryotic vs. prokaryotic metacommunity structure, the microeukaryotic community composition of 15 marine-derived saline lakes, one freshwater lake as well as one coastal marine site in the Vestfold Hills were analyzed using high-throughput DNA sequencing of the V4 region of the 18S rRNA gene. Comparable data from prokaryotes was obtained from Logares et al. (2013) [16S rRNA gene, regions V3-V4]. We hypothesized that, similar to prokaryotes (Logares *et al*. 2013), microeukaryotes would mostly be structured by environmental selection associated to the steep salinity gradient. In contrast, we found that microeukaryotes were structured mostly by ecological drift, with a minor role for salinity-driven environmental selection.

## MATERIALS AND METHODS

### Field sampling, 18S rRNA-gene sequencing, and bioinformatics analysis

Our field sampling area is located in the Vestfold Hills, Eastern Antarctica (68°S, 78°E, Fig. S1). Water samples from different depths were collected in 16 selected lakes as well as in one coastal marine site (Fig. S1, Table S1) during the austral summer (December 2008-February 2009). Sample salinity, nutrients (P = phosphate, N = nitrate), dissolved organic carbon (DOC), depth, water temperature, lake area and altitude were measured (Table S1). These variables were standardized as z-scores, that is, deviations of the raw values from the mean in standard deviation units. Furthermore, salinity was divided into categories following Logares et al. (2013): freshwater (0 PSU), low-brackish (1-6 PSU), high-brackish (7-30 PSU), marine salinity (31-36 PSU), hypersaline (43-75 PSU), and high-hypersaline (76-250 PSU) (Table S1). Microbes were collected onto polycarbonate filters (Supor-200, 47mm; PALL Corporation, East Hills, NY, USA) and subsequently filters were stored at −80°C. Community DNA was extracted from the filters and the hypervariable V4 region of the 18S rRNA gene was PCR-amplified and sequenced in a Roche 454 platform (GS FLX Titanium) as explained in Supporting Information. The produced sequences (thereafter “reads”) were processed with QIIME v1.3.0 (Caporaso *et al*. 2010) generating Operational Taxonomic Units (OTUs; species proxies) with a 99% similarity threshold (see Supporting Information for a detailed description of the analyses). Sequence data was deposited in the European Nucleotide Archive (ENA) public database with the references ERS1806710-37.

In microeukaryotes, our initial full OTU table consisted of 991 OTUs (excluding Metazoa), including 666,151 reads distributed across 25 samples (Table S1). A second OTU table was generated excluding singletons, which contained 763 OTUs and 665,923 reads. Then, a third subsampled OTU table was generated using rrarefy in Vegan (Oksanen *et al*. 2008), consisting of 420 OTUs and 1,273 reads per sample (total 31,825 reads). For prokaryotes, we have used as initial OTU tables those indicated in Logares et al. (2013).

In order to compare microeukaryotes and prokaryotes as well as to construct association networks, we have generated a specific dataset with the same samples available for both microeukaryotes and prokaryotes (given that not all samples available for microeukaryotes had their prokaryotic counterpart available or vice versa). This dataset included 20 samples (Table S1), with 378 OTUs in microeukaryotes and 1,192 OTUs in prokaryotes.

Regionally rare or abundant OTUs were arbitrarily defined as those with mean relative abundances <0.01% or >0.1% respectively (see Logares *et al*. 2014).

### Community ecology and phylogenetic analyses

Most analyses were performed in R (R-Development-Core-Team 2008) using the package Vegan (Oksanen *et al*. 2008). Beta diversity was estimated using Bray-Curtis dissimilarities as well as gUnifrac (Chen *et al*. 2012), based on Lozupone and Knight (2005). For gUnifrac, as well as for determining Faith’s Phylogenetic Diversity (PD) [Faith 1992] and Mean Nearest Taxon Distance (MNTD) [Webb *et al*. 2002], we constructed phylogenies based on the reads representing each OTU. Reads were aligned against an aligned reference database (SILVA v108) using Mothur (Schloss *et al*. 2009). RAxML (Stamatakis *et al*. 2005) was used to infer 100 phylogenetic trees under the GTR+CAT/G+I model, and the best topology was selected. PD and MNTD were inferred using Picante (Kembel *et al*. 2010); MNTD values were compared to a null model that shuffled taxa-labels 1,000 times across all taxa included in the phylogeny-based distance matrix (Kembel 2009).

PERMANOVA analyses were used to investigate what proportion of community variance was explained by the selected environmental variables. Furthermore, we analyzed the correlation between environmental variables and community differentiation using Mantel and Partial Mantel tests (Legendre & Legendre 1998) based on Pearson correlations between Bray Curtis (community data) and Euclidean (standardized environmental data) dissimilarities. Mantel tests were run with 999 permutations in Vegan. Furthermore, Mantel tests as well as Mantel correlograms (Oden & Sokal 1986) were used to explore the relationships between community dissimilarity and geographic distance.

In order to explore whether deterministic vs. stochastic processes influenced community assembly we calculated the Raup-Crick metric (Chase *et al*. 2011) using Bray-Curtis dissimilarities that consider relative abundances [RC_bray_], as proposed by Stegen *et al*. (2013). This metric compares the measured β-diversity against the β-diversity that would be obtained under random community assembly by using a null distribution derived from a randomization. For each pair of communities, the randomization was run 999 times, generating a null distribution of Bray-Curtis dissimilarities (Stegen *et al*. 2013). Only OTUs with >100 reads were included, as those have high chances to be part of the regional species pool (Chase *et al*. 2011). RC_bray_ values > +0.95 or < −0.95 are interpreted as significant departures from a stochastic community assembly, pointing to environmental selection or dispersal limitation. On the contrary, RC_bray_ values between −0.95 and +0.95 point to a stochastic community assembly (Chase *et al*. 2011).

### Habitat specialists or generalists

Habitat (salinity) specialization was determined using Levins’ niche breadth (Levins 1968) and the six mentioned environmental types (Freshwater, Low-Brackish, High Brackish, Marine salinity, Hypersaline, and High-Hypersaline). Habitat generalists were defined as those OTUs having a niche breadth index *B*≥3. Since low-abundance taxa may be temporarily absent or below detection limits in some environments, they could erroneously appear as specialists when using Levin’s niche breadth. For this reason, to determine specialist taxa, we only considered OTUs with >50 reads, being present in a maximum of two environmental categories and with a *B* value that was significantly lower than a random distribution. Randomizations of *B* were run with the R package EcolUtils (Salazar 2015). To further investigate specialist taxa restricted to only one environmental type we used the INDicator VALues analysis (Dufrene & Legendre 1997) as implemented in the R package labdsv (Roberts 2016).

### Association networks analyses

We constructed association networks in order to explore whether groups of OTUs with similar salinity preferences populate different lakes. Only microeukaryotic or prokaryotic OTUs with a minimum of 50 reads and which were present in at least two samples were analyzed (total of 156 OTUs). Association networks were generated using Conet (Faust *et al*. 2012), and positive as well as negative correlations were inferred with 5 methods: Bray-Curtis, Kullback-Leibler, Mutual information, Pearson, and Spearman. P-values were calculated through permutations and bootstrapping, considering only significant (p<0.05) correlations. Networks were visualized in Cytoscape (Shannon *et al*. 2003). Highly interconnected regions of the network or clusters were detected using MCODE (Bader & Hogue 2003), considering co-occurrences only.

## RESULTS

### Structuring patterns in microeukaryotic vs. prokaryotic communities

Bray-Curtis dissimilarities between microeukaryotes and prokaryotes occurring in the same community were weakly correlated (Mantel test *ρ*=0.28, p=0.003; Fig. 1A), indicating that both domains have different patterns of community structure. Furthermore, the analysis of the Raup-Crick metric (RC_bray_) indicated that most (85%) RC_bray_ values for prokaryotes are above +0.95 or below −0.95, pointing to environmental selection or dispersal limitation as having a role in community assembly (Fig. 1B). In contrast, most (71%) microeukaryotes presented RC_bray_ values between −0.95 and +0.95, indicating that stochastic processes (drift) have been predominant during the assembly of their communities. Congruently, correlations between microeukaryotic beta diversity and salinity, phosphorous, nitrogen and DOC were weak or non-significant. Microeukaryotic beta diversity presented a weak positive correlation with salinity (Mantel *ρ* = 0.30, p=0.003; Figs. S2-S3). That correlation did not change substantially when controlled by phosphorous (partial mantel *ρ*=0.24, p=0.02) or temperature (partial mantel *ρ*=0.22, p=0.03), but it became non-significant when controlled by nitrogen (partial mantel p=0.31) or DOC (partial mantel p=0.09). The correlation between phosphorous and microeukaryotic community composition was significant when controlled by salinity (partial mantel *ρ*=0.26, p=0.03). Yet, correlations between microeukaryotic community composition and temperature, nitrogen or DOC were not significant when controlled by salinity (partial mantel: temperature, p=0.23; nitrogen, p=0.33; DOC, p=0.39). Additional analyses indicated that salinity explained about 8% of microeukaryotic community variance (PERMANOVA, p=0.01). Salinity categories explained 9.2% of microeukaryotic phylogenetic (gUnifrac) variance (Permanova, p=0.01), but Mantel tests did not indicate significant correlations between gUnifrac distances and salinity change (Euclidean distances) [p=0.13]. Interestingly, lake altitude (that is, lake surface water level in relation to the sea level), which can be related to some extent with lake age in this system, was weakly correlated with microeukaryotic community composition (mantel *ρ*=0.19, p=0.04), although this correlation became non-significant when controlling by salinity (partial mantel p=0.54). Other variables that were not correlated with microeukaryotic beta diversity were oxygen (p=0.32), lake area (p=0.52), volume of sampled water (p=0.38) and lake maximum depth (p=0.47).

**Figure 1.**
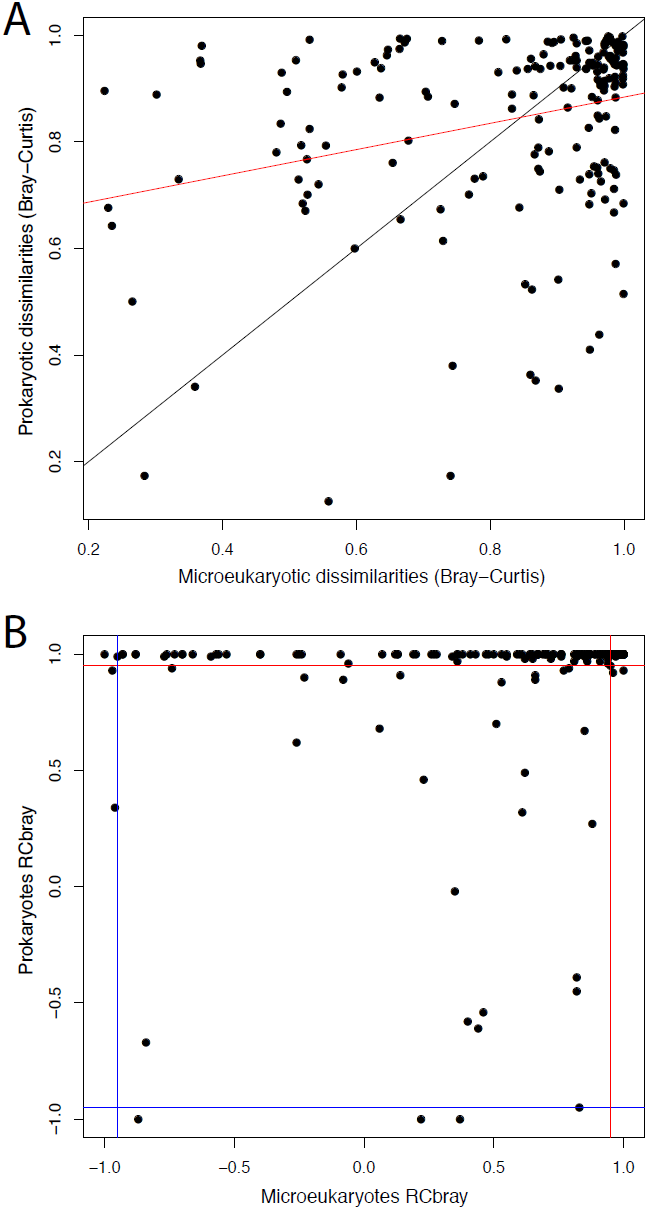
Panel A. Microeukaryotic vs. prokaryotic Bray-Curtis dissimilarities. The black line indicates the 1:1 line, while the red line shows the best linear fit (R^^2^^ = 0.07, slope = 0.24; p<0.05). Mantel test *ρ* =0.28, p=0.003. Panel B. Raup-Crick values between microeukaryotic and prokaryotic communities from the same samples. Values of RC_bray_ greater than +0.95 or less than −0.95 indicate that environmental selection or dispersal limitation influence community structuring. On the contrary, RC_bray_ values between −0.95 and +0.95 point to stochasticity during community assembly. Blue and red lines indicate −0.95 and +0.95 RC_bray_ values respectively. Note that most (85%) RC_bray_ values for prokaryotes are above +0.95 or below −0.95, pointing to environmental selection or dispersal limitation having a role in their community assembly. In contrast, most (71%) microeukaryotes present RC_bray_ values between −0.95 and +0.95, indicating that stochastic processes predominate during the assembly of their communities.

In both microeukaryotes and prokaryotes, the regional abundance of OTUs (that is, the mean abundance of OTUs across communities) was positively correlated with the number of samples in which OTUs were present (i.e. frequency of occurrence) [Fig. 2 A - B]. In both domains, the proportion of OTUs that occurred in >50% of the samples was similar (1.4% in microeukaryotes and 1.6% in prokaryotes). Among microeukaryotes (Fig. 2 A), 20.1% of the OTUs (76 OTUs) were regionally abundant while 41.0% of the OTUs (155 OTUs) were regionally rare. In prokaryotes (Fig. 2 B), regionally abundant OTUs represented 8.5% of OTUs (102 OTUs), while 66% of the OTUs (787 OTUs) were regionally rare. In microeukaryotes, beta diversity between abundant vs. rare taxa showed a weak but significant positive correlation (mantel *ρ*=0.27, p=0.001). Both microeukaryotes and prokaryotes showed a weak-moderate positive correlation between community differentiation and geographic distance (isolation by distance) within the first 400m (Fig. 2 C - D). This correlation was higher in prokaryotes (Fig. 2 D).

**Figure 2.**
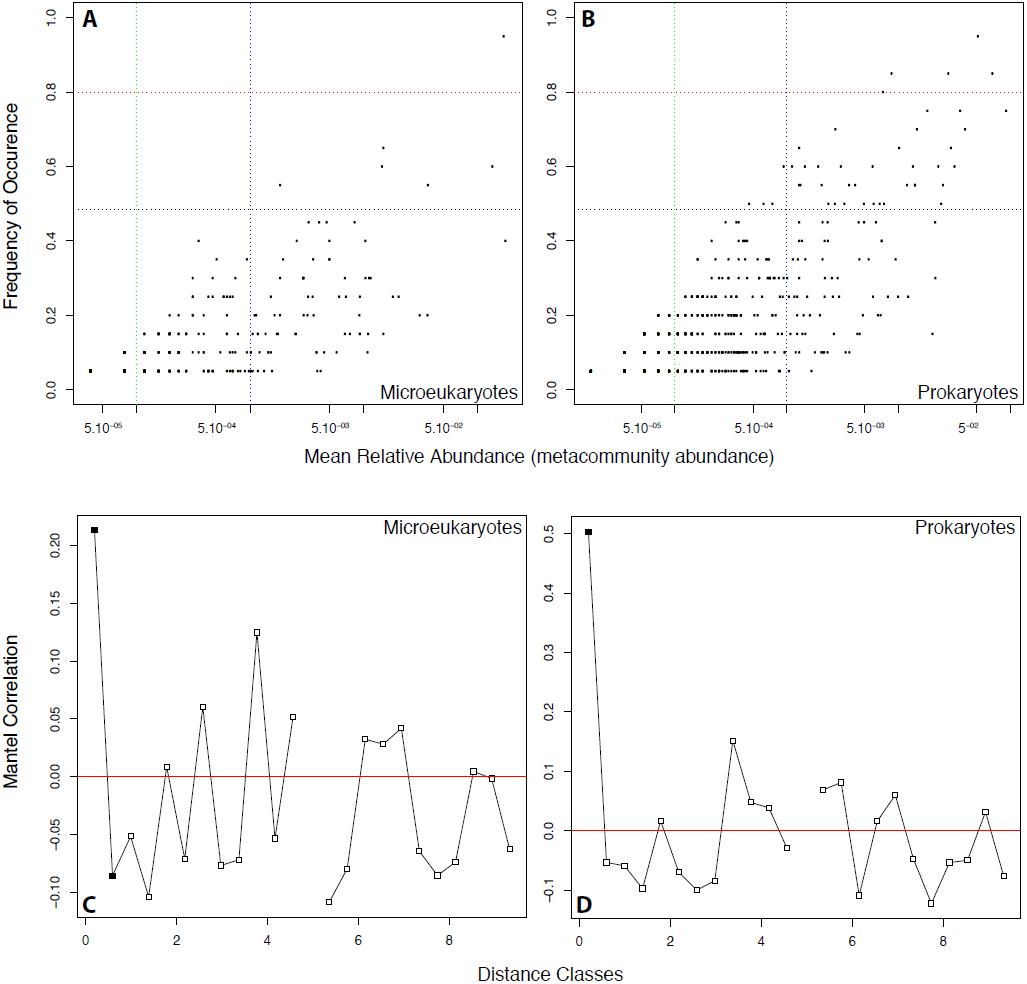
Panels A and B: OTU mean relative abundance vs. occurrence. Each dot is an OTU. The red and black horizontal lines separate OTUs that occur in more than 80% and 50% of the samples respectively. A total of 1.6% (43 out of 2586 OTUs) of prokaryotes and 1.4% (6/420) of microeukaryotes were present in >50% of the samples. In both panels, the blue and green vertical lines indicate regionally abundant (mean relative abundance >0.1%) or rare (mean relative abundance <0.01%) OTUs respectively. In microeukaryotes (Panel A), 20.1% of the OTUs (76 OTUs) were regionally abundant while 41.0% of the OTUs (155 OTUs) were regionally rare. In prokaryotes (Panel B), regionally abundant OTUs represented 8.5% of OTUs (102 OTUs), while 66% of the OTUs (787 OTUs) were regionally rare. Panels C and D indicate Mantel correlograms for microeukaryotes and prokaryotes respectively. The ordinate indicates mantel correlations (positive or negative) between geographic distance and community dissimilarity while the abscissa indicates geographic distance classes. Black squares indicate significant (p<.05) correlation values, while white squares indicate non-significant correlations. Distance classes have about 400m. Classes for which data are not available (“NA”) are not shown.

Local microeukaryotic richness did not approach saturation in most samples (Fig. S4). On the other hand, regional richness seems to have been better recovered, with a saturation curve starting to level off (Fig. S5), a pattern that was congruent with accumulation curves (Fig. S6). Microeukaryotic diversity (Shannon index; Fig. S7) and Faith’s Phylogenetic Diversity (PD; Fig. S8) did not show well-defined trends with salinity. Microeukaryotic taxonomic diversity is indicated in Supporting Information and Fig. S9, while prokaryotic diversity is presented in Logares et al. (2013).

Microeukaryotic beta diversity displayed a moderate positive correlation with Unifrac phylogenetic distances (Mantel test *ρ*=0.50, p=0.001; Fig. S10), indicating that community differentiation based on OTUs is mirrored to some extent by phylogeny. Mean Nearest Taxon Distances (MNTD) were analyzed in order to determine whether microeukaryotic communities showed phylogenetic over-clustering or over-dispersion patterns (Webb *et al*. 2002, Cavender-Bares *et al*. 2009). When MNTD was not weighted by OTU abundances, a total of 14 out of 25 (56%) communities showed phylogenetic over-clustering, while none of them showed over-dispersion (Table S2). When considering OTU abundances, 8 out of 25 (32%) of the microeukaryotic communities presented phylogenetic over-clustering while none of them presented over-dispersion (Table S3).

### Microeukaryotes across the marine-lacustrine boundary

A relatively few, but regionally abundant microeukaryotic OTUs were shared between the lakes and the sea (Fig. 3 A). Considering OTUs with >25 reads, only 32.1% of them (101 OTUs) were shared between lakes and the sea, while 65.7% (207 OTUs) were exclusive to lakes or to the sea (2.2%; 7 OTUs) [Fig. 3 A]. The regional abundance of OTUs occurring in lakes (mean abundances across 24 samples) was compared to the corresponding abundance in the sea. We found that OTUs shared between the lakes and the sea were the most abundant (78.5% of reads [total 41,256 reads]), while OTUs restricted to lakes represented 20.7% (10,891 reads) of the total abundance and those restricted to the sea represented 0.8% (424 reads) [Fig. 3 B].

**Figure 3.**
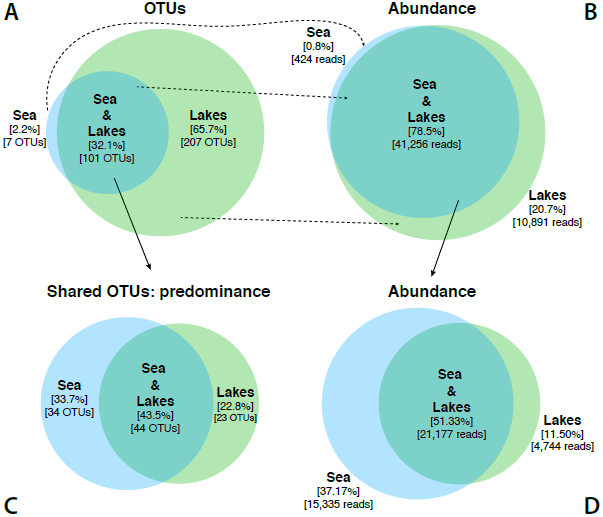
Venn diagrams indicating A) the number of OTUs that were exclusive to the lakes and the sea, as well as the OTUs shared between these environments. Panel B) shows the corresponding abundance (as number of reads) of the OTUs shown in panel A) [indicated with dashed arrows]. OTUs that were detected in both marine and lacustrine environments were further classified as predominantly marine or lacustrine depending on whether their abundances presented a difference (ratio) >10 between environments (solid arrows). For the latter OTUs, Panel C shows habitat predominance, while panel D shows the corresponding abundance. Only OTUs with >25 reads in the full OTU table were considered.

OTUs that were present in both the lakes and the sea were not necessarily evenly distributed across these environments. These OTUs were classified as predominantly marine or lacustrine depending on whether their abundances presented a ratio >10 between environments. Otherwise, OTUs were regarded as “even”. A total of 44 OTUs out of the shared 101 OTUs were evenly distributed between marine and lacustrine environments, while 23 OTUs were predominantly lacustrine and 34 OTUs were predominantly marine [Fig. 3 C]. OTUs that were more or less evenly distributed across the lakes and the sea were the most abundant (51.33%), while predominantly marine or lacustrine OTUs accounted for 37.17% and 11.50% of the abundance respectively [Fig. 3 D].

### Specialist and generalist microeukaryotes

We calculated what proportion of OTUs could be regarded as specialists or generalists in relation to salinity. For that, habitats were classified as Freshwater, Low-Brackish, High Brackish, Marine salinity, Hypersaline, and High-Hypersaline following Logares et al. (2013). Levins’ niche breadth (*B)* [Levins 1968] was calculated in relation to these habitats and the obtained values ranged between 1 and 3.9, considering a theoretical maximum *B* of 6 (number of salinity habitats) [Fig. S11]. Obtained values were contrasted against a null model (randomization), in which *B* ranged between 3 and 1. Considering the randomized values, we determined that OTUs with *B*≥3 had good chances to be habitat generalists. In addition, we performed a second randomization using incidence data only, which served to detect habitat specialists. After running these analyses, only OTUs with >50 reads were retained (Table 1). Altogether, these analyses indicated 6 microeukaryotic OTUs (out of a total of 259) that can be regarded as habitat generalists and 5 that can be considered specialists (Table 1). To detect strict habitat specialists (that is, OTUs specialized in a single environment), we used the Dufrene and Legendre (1997) index for detection of indicator species. A total of 3 OTUs (>25 reads each) associated to fresh waters were indicated as strict specialists (Table S4).

**Table 1.** Habitat generalists or specialists microeukaryotes

| OTUs | Ab [%]* | B† | Type‡ | Fr§ | LB§ | HB§ | Ma§ | Hy§ | HH§ | Acc.#¶ | % Similarity** | Taxonomy†† |
| --- | --- | --- | --- | --- | --- | --- | --- | --- | --- | --- | --- | --- |
| 1094 | 19.35 | 3.83 | G | 676 | 332 | 332 | 310 | 118 | 65 | EF417318 | 97.5 | Dinophyceae; *A. aff. malmogiense* (Mc-Neil-A) |
| 330 | 17.82 | 3.95 | G | 23 | 356 | 192 | 292 | 176 | 14 | EF432522 | 98.0 | Cryptophyceae; Cryptophyta; *Geminigera* |
| 620 | 1.37 | 3.16 | G | 0 | 6 | 42 | 18 | 28 | 2 | HQ111509 | 98.0 | Chlorophyceae; *Pyramimonas gelidicola* |
| 1022 | 1.30 | 3.28 | G | 35 | 30 | 23 | 0 | 0 | 6 | EF417318 | 95.7 | Dinophyceae; *A. aff. malmogiense* (Mc-Neil-A) |
| 1437 | 0.96 | 3.73 | G | 13 | 11 | 34 | 12 | 2 | 6 | JF826331 | 96.8 | Alveolata |
| 847 | 0.90 | 3.18 | G | 1 | 9 | 12 | 38 | 11 | 5 | GU647176 | 98.5 | Dinophyceae; *Polarella glacialis* |
| 901 | 4.13 | 1.59 | S | 0 | 0 | 0 | 0 | 86 | 264 | KC487270 | 96.9 | Chlorophyceae; *Dunaliella* |
| 1373 | 2.51 | 1.00 | S | 0 | 0 | 199 | 0 | 0 | 0 | JF343797 | 95.0 | Chlorophyceae; *Nannochloris* |
| 785 | 1.20 | 1.00 | S | 0 | 0 | 95 | 0 | 0 | 0 | JF343797 | 92.1 | Chlorophyceae; *Nannochloris* |
| 327 | 0.55 | 1.00 | S | 172 | 0 | 0 | 0 | 0 | 0 | GU117581 | 97.5 | Chlorophyceae; *Chlamydomonas* |
| 550 | 0.47 | 1.00 | S | 0 | 0 | 0 | 73 | 0 | 0 | EF140624 | 98.3 | Bacillariophyceae; *Fragilariopsis* |

Habitat generalists were defined as those having a niche breadth index $B \geq 3$ , while specialists (shaded area) were defined as those being present in maximum 2 environmental types with a distribution that differs significantly from chance. All considered OTUs presented >50 reads.
\* Total abundance in the metacommunity of each OTU as percentage (i.e. percentage of total reads).
† Niche breadth $B$ values.
‡ G=habitat generalists, S=habitat specialist.
§ Number of reads. Fr=freshwater, LB=low-brackish, HB=high brackish, Ma=marine salinity, Hy=hypersaline, HH=high-hypersaline.
¶ Genbank accession numbers of the best blast hits.
\*\* Percentage of similarity of the best blast hit (coverage >80%).
†† Taxonomic assignment of the best blast hit according to SILVA v119.1.

### Differential response to salinity in microeukaryotes

We used a correlation network approach to determine which groups of OTUs exhibit differential response to salinity. The clustering of the network containing only positive edges indicated 4 clusters containing >7 nodes (OTUs) that included microeukaryotes and prokaryotes, and which were further analyzed (Table 2; Supporting information). Most generated clusters and sub-clusters could be associated to salinity categories (Table 2) or sites, pointing to OTUs with differential salinity preference and, in some cases, habitat restriction. Cluster 1 (Table 2 & S5; Fig. S12) includes the Hypersaline OTU sub-cluster (1a) as well as the High-Hypersaline sub-cluster (1b). Cluster 2 was restricted to High-Brackish waters [Table 2 & S5; Fig. S13]. Cluster 3 was more diverse in terms of the salinity of the included samples, and could be subdivided into 6 sub-clusters (Fig. S14; Table 2 & S5), which included samples ranging from freshwater to marine salinity. Cluster 4 included the sub-cluster 4b containing High-Brackish to Hypersaline samples (Table 2 & S5; Fig. S15). In summary, network analyses showed evidence of a total of 30 OTUs with differential response to salinity (Table S5), which represented 63.17% of the total OTU abundance.

**Table 2.** Network clusters

| Cluster* | Score† | Nodes‡ | Edges§ | Domain¶<br>(B-E) | # Lakes** | Marine<br>presence†† | Distribution salinity categories‡‡ | Preferred salinity§§ |
| --- | --- | --- | --- | --- | --- | --- | --- | --- |
| **1** | 11.6 | 24 | 134 | - | 3 | - | - |  |
| 1a | - | - | - | B(9)+E(5) | 1 | No | Hy(100.0; 100.0) | **Hy** |
| 1b | - | - | - | B(8)+E(2) | 2 | No | Hy (50.0; 16.4), HH(50.0;83.6 ) | **HH** |
| **2** | 7.0 | 7 | 21 | B(5)+E(2) | 1 | Yes | HB(50.0; 88.3), Ma(50.0; 11.7) | **HB** |
| **3** | 6.7 | 41 | 134 | - | - | - | - |  |
| 3a | - | - | - | B(5)+E(4) | 1 | Yes | Ma(100.0; 100.0) | **Ma** |
| 3b | - | - | - | B(7)+E(1) | 3 | No | Fr(20.0; 80.3), LB(60.0; 14.1), HB(20.0; 5.6) | **Fr** |
| 3c | - | - | - | E(8) | 6 | No | Fr(14.3; 14.2),LB(42.8; 51.5), HB(28.6; 29.3),<br>Ma(14.3; 5.0) | **LB+HB** |
| 3d | - | - | - | B(3)+E(2) | 2 | No | LB(66.7; 77.7), HB(33.3; 22.3) | **LB+HB** |
| 3e | - | - | - | B(5)+E(2) | 13 | No | LB(50.0; 45.8), HB(18.7; 25.0), Ma(6.3;<br>19.0), Hy(25.0; 10.2) | **LB+HB+Ma** |
| 3f | - | - | - | B(2)+E(2) | 2 | No | LB(100.0; 100.0) | **LB** |
| **4** | 6.5 | 17 | 52 | - | - | - | - |  |
| 4a | - | - | - | B(9) | 6 | No | LB(50.0;41.0), HB(50.0; 59.0) | **LB+HB** |
| 4b | - | - | - | B(6)+E(2) | 3 | No | HB(50.0; 21.4), Ma(1; 23.0), Hy(50.0; 55.6) | **HB+Ma+Hy** |
| **5** | 4.0 | 4 | 6 | - | - | - | - |  |
| **6** | 3.3 | 4 | 5 | - | - | - | - |  |
| **7** | 4.0 | 4 | 6 | - | - | - | - |  |
| **8** | 3.0 | 5 | 3 | - | - | - | - |  |

\* Cluster number.
† Highest scoring node in the cluster.
‡§ Number nodes and edges in the cluster.
¶ Number of OTUs belonging to different domains present in the cluster (B=Bacteria, E=Eukaryota).
\*\* Number of lakes in which members of the cluster are present.
†† Presence or absence of members of the cluster in coastal marine waters.
‡‡ Salinity categories of the samples where OTUs belonging to different clusters occur; The data is presented as “Salinity Category (percentage of samples within the category; corresponding percentage of reads)” [codes: Fr=freshwater, LB=low-brackish, HB=high brackish, Ma=marine salinity, Hy=hypersaline, HH=high-hypersaline].
§§ Salinity category of the samples which represent >80% of the reads in each cluster

### Contrasting distribution patterns for abundant OTUs of the same taxon

We found distinctive OTU-distribution patterns for different abundant OTUs of *A.* aff. *malmogiense* (9 OTUs with >2,000 reads), pointing to historical effects. Overall, OTUs 1094, 1022, 830, 286 & 375 were prevalent in Lake Abraxas, Ace Lake, Crooked Lake, Ekho Lake, Lake Hand & Highway Lake, while OTUs 305, 54, 1064 & 1197 were predominant in Lake McNeil and Vereteno Lake (Fig. 4). A phylogeny of these OTUs indicated that sequence divergence did not follow the same patterns as the OTU distribution between lakes (Fig. S16).

**Figure 4.**
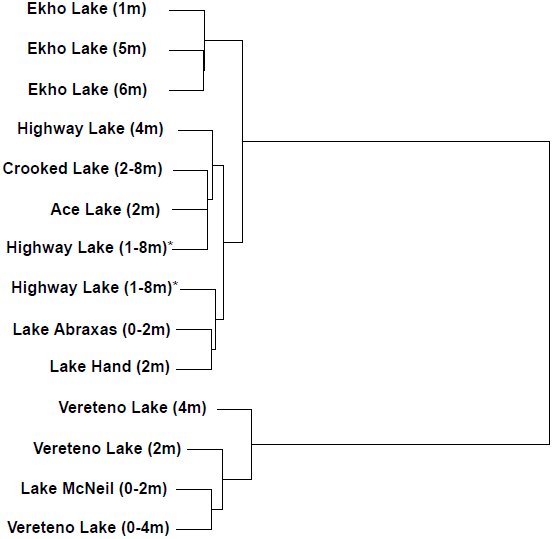
UPGMA dendrogram showing sample similarity in terms of their composition of *Apocalathium* aff. *malmogiense* OTUs (9 OTUs total). Both OTUs and samples with >2,000 reads were used. Each dendrogram leaf corresponds to one sample, and the depth from where it was taken is indicated within parentheses. *Different samples from Highway Lake integrating the water column.

## DISCUSSION

### Microeukaryotes are structured by ecological drift while prokaryotes by selection

Previous studies on the prokaryotic microbiota occurring in the same set of lakes indicated that the main structuring force was environmental selection due to salinity [Mantel *ρ* = 0.7, p=0.001; correlation between salinity and community differentiation] (Logares *et al*. 2013). Naturally, we hypothesized that the microeukaryotic community would be similarly structured by salinity. However, our results indicated that salinity [Mantel *ρ* = 0.3, p=0.003], as well as the other measured environmental variables, have a weak effect on microeukaryotic community structure. In addition, our analyses using the Raup-Crick metric indicated that while 85% of community turnover in prokaryotes could be associated to environmental selection or dispersal limitation, 71% of community turnover in microeukaryotes could be related to stochastic processes (ecological drift). In the case of prokaryotes, given that salinity could explain a considerable fraction of community dissimilarity, and that we have selected OTUs which display a moderate or high regional abundance, we propose that this environmental factor, instead of dispersal limitation, is deterministically driving community turnover. Very few studies have investigated the structure of prokaryotic vs. microeukaryotic communities. In one study carried out in the Baltic Sea, considering salinities between 2 and 22, it was found that beta diversity of both microeukaryotes and prokaryotes were correlated across the salinity gradient (Hu *et al*. 2016). This suggests that environmental selection driven by salinity had similar effects in both prokaryotes and microeukaryotes. In another study, contemplating salinities between 0-30 in a river to ocean transect, it has been found that prokaryotes were more broadly distributed across the salinity gradient than microeukaryotes, which tended to be found in narrower salinity ranges (Nawar 2016). Here, it appears that salinity change is affecting differently the community structure of both domains. The latter two studies contrast with our results, but it should be taken into account that the salinity gradient we studied is about 10 times larger (0-250) than the mentioned studies, and that we investigated lakes instead of more connected habitats, such as rivers or the sea. In addition, local adaptation to multiple salinities could have expanded the salinity range of some species inhabiting the studied lakes. There is evidence of local adaptation to a wider salinity range in strains of the dinoflagellate *Polarella glacialis* living in some of the studied saline lakes, when compared to marine ancestors (Rengefors *et al*. 2015). Furthermore, and more in line with our results, a recent work reports that deterministic processes governed community turnover in soil bacteria while community turnover of soil fungi (yeasts and hyphae-forming variants) from the same samples was mostly influenced by stochastic processes (Powell *et al*. 2015). Thus, it appears that microeukaryotes and prokaryotes could display contrasting or similar community structuring patterns depending on the characteristics and history of different habitats.

The weak covariation we found between community and habitat dissimilarities in microeukaryotes could also be partially explained by the presence of a few abundant habitat generalists (that is, halotolerant taxa) in the metacommunity that were detected along the salinity gradient. Furthermore, we found relatively few habitat (salinity) specialist OTUs in microeukaryotes (5 defined with niche breadth *B* as well as 3 strict specialists defined with IndVal). A higher number of specialists would have been expected in a scenario of strong environmental selection. This contrasts markedly with the specialist prokaryotic OTUs reported for the same lakes (939 defined using *B* and 365 defined using IndVal; Logares *et al*. 2013). The latter agrees with other studies indicating a dominance of specialist OTUs in diverse prokaryotic communities (Mariadassou *et al*. 2015).

The limited effects of environmental selection in microeukaryotic communities were also reflected, to some extent, by phylogeny (using the Mean Nearest Taxon Distances [MNTD]). In MNTD analyses, over-clustering suggests environmental selection and over-dispersion patterns point to competition (Webb *et al*. 2002, Cavender-Bares *et al*. 2009). We found that 32% of the communities presented over-clustering (environmental selection) when OTU abundances were considered, while there was no evidence of over-dispersion (competition).

We have also found that in contrast to prokaryotes, abundant vs. rare microeukaryotic sub-communities displayed different distribution patterns. Beta diversity in regionally abundant (mean relative abundance >0.1%) vs. rare (mean relative abundance <0.01%) microeukaryotic sub-communities showed only a weak positive correlation (mantel *ρ*=0.27, p=0.001) pointing to decoupled structuring patterns. In contrast, prokaryotes showed a higher correlation between beta diversity associated to abundant vs. rare taxa (mantel *ρ*=0.82, p<0.01; Logares *et al*. 2013). Thus, it appears that while in prokaryotes both abundant and rare members of communities were subject to the same structuring forces (environmental selection), in microeukaryotes, there were either different processes structuring these sub-communities (including historical processes) or differential dispersal limitation. Abundant or rare sub-communities of marine microeukaryotes from several coastal locations of Europe also presented different structuring patterns (Logares *et al*. 2014) suggesting that these components may normally be decoupled in this domain.

### Taxon-specific responses to salinity and historical effects

The absolute number of salinity specialists was two orders of magnitude lower in microeukaryotes than in prokaryotes (Logares *et al*. 2013), suggesting that the former have, in general, wider niche breadths and possibly a less deterministic response to salinity compared to prokaryotes. Accordingly, we found that whole microeukaryotic communities were only weakly correlated with salinity or other environmental parameters. Yet, there were microeukaryotic OTUs that displayed a distribution correlated to salinity (as indicated by network analyses), pointing to environmental selection. Thus, it appears that while some members of the microeukaryotic community are structured by environmental selection, most of the others are structured by more stochastic processes. Comparable scenarios were found in other studies in prokaryotes. For example, it has been found that during the assembly of prokaryotic communities, both environmental selection as well as neutral processes can have a role (Langenheder & Szekely 2011). Similarly, Lindh et al. (2016) found that different taxonomic groups of bacterioplankton in the Baltic Sea were structured by environmental filtering, mass effects, patch dynamics or the neutral model (see Leibold *et al*. 2004).

Overall, both predictable and unpredictable factors seem to have a role in the assembly of most communities (Chase 2007). In cases where communities could develop multiple states or equilibria under similar environmental conditions, then the resulting community configuration may reflect assembly history (Chase 2003). Furthermore, historical processes or priority effects may be more prevalent in environments exerting a lower selective pressure (Chase 2007). Thus, it could be hypothesized that non-extreme salinity concentrations may allow for historical effects. Our results indicating a heterogeneous distribution for 9 abundant OTUs of *A.* aff. *malmogiense* among lakes, some physicochemically highly similar, support the latter hypothesis. In addition, the latter suggests that historical processes may have had a higher prevalence in microeukaryotes than in prokaryotes, given that the former were predominantly decoupled from salinity variation.

For both microeukaryotes and prokaryotes, we observed weak or moderate positive correlations of geographic distance with community dissimilarity within the first 400m. The overall shape of the correlogram could be related to a “single bump” spatial structure (Legendre & Legendre 1998), which points to a loss of spatial autocorrelations after 400m. Within these 400m, our results indicated a more defined distance-decay pattern in prokaryotes than in microeukaryotes, which could be linked to a higher dispersal of at least some microeukaryotic OTUs compared to prokaryotes. Nevertheless, the latter results point to limited effects of contemporary dispersal on community composition within 400m. This agrees with studies in dinoflagellates populating lakes in the same area for which there was evidence of limited gene flow due to potential dispersal limitation (Rengefors *et al*. 2012). A limited dispersal could contribute to historical effects, especially in microeukaryotes. Yet, the amount of dispersal occurring in the investigated area could be enough for ensuring environmental selection in prokaryotes.

### Regional and global dispersal

Most microeukaryotic OTUs detected in the sea were also present in lakes, pointing to a widespread presence of the coastal marine microbiota in lakes. Only a small percentage of microeukaryotic OTUs [2.2%, 7 OTUs] were detected as being exclusive to the sea. Similarly, in prokaryotes, only a small percentage of OTUs was exclusive to the sea [6.7%, 173 OTUs] (Logares *et al*. 2013). Overall, microeukaryotic OTUs which were present in both lakes and the sea represented 32.1% of the total OTUs [101 OTUs], which accounted for 78.5% of the total OTU abundance. In prokaryotes, 6.5% of the total OTUs (167 OTUs) were shared between lakes and the sea, representing 45.5% of the total abundance (Logares *et al*. 2013). A total of 56.4% of the microeukaryotic OTUs present in both marine and lacustrine environments (representing 48.7% of the shared lacustrine-marine abundance) presented a difference of >10 times in abundance between habitats, suggesting that the presence in the lakes of some marine OTUs (and vice versa) was due to immigration from the neighboring sea, but that such OTUs will not establish viable populations (mass effects). In contrast, those OTUs which are part of established populations in both the sea and lakes could be associated to recent - ancient dispersal or they could partially represent the original marine microbiome that populated the marine bays that were isolated from the sea and became lakes.

In total, 65.7% of the microeukaryotic OTUs (20.7% of the total reads [total reads = averaged lacustrine reads plus marine reads]) were restricted to lakes. In prokaryotes, 86.8 % of the OTUs (54% of the reads) were restricted to lakes (Logares *et al*. 2013). These lake-restricted taxa could be the product of global or regional dispersal (Montresor *et al*. 2003; Pearce *et al*. 2007; Logares *et al*. 2008; Ghiglione *et al*. 2012; Logares *et al*. 2013), or represent Antarctic endemic taxa (Vyverman *et al*. 2010). In sum, the investigated habitats (lakes) appear to have received a considerable amount of regional or global immigrants during their diversification, which may have promoted a larger amount of historical or priority effects in microeukaryotes than in prokaryotes.

## Conclusion

Habitat diversification can promote contrasting community structuring mechanisms in planktonic microeukaryotes and prokaryotes. Therefore, we consider both community components should be analysed together in future ecological studies in order to attain a comprehensive understanding of the processes that shape microbial assemblages.

## ACKNOWLEDGEMENTS

This project was supported by a Linnaeus grant to the Centre for Animal Movement Research (CAnMove, Lund University; Swedish Research Council grant number 349-2007-8690). In addition, the project was supported with a grant from the Swedish Research Council (621-2012-3726) to KR. RL was supported by a Ramón y Cajal fellowship (RYC-2013-12554, MINECO, Spain). Sample collection was supported by grant from the Australian Antarctic Research Assessment Committee awarded to KR and Johanna Laybourn-Parry. We thank Johanna Laybourn-Parry as well as the Australian Antarctic Division staff in Kingston, onboard the research vessel Aurora Australis and at Davis Station, Antarctica. High-Performance computing analyses were run at the MARBITS bioinformatics platform of the Institut de Ciències del Mar in Barcelona, Spain.

